# Genome-wide identification of long non-coding RNAs in the gravid ectoparasite *Varroa destructor*

**DOI:** 10.1101/2020.06.14.151340

**Authors:** Zheguang Lin, Yibing Liu, Xiaomei Chen, Cong Han, Wei Wang, Yalu Ke, Xiaoling Su, Heng Chen, Hao Xu, Guohong Chen, Ting Ji

## Abstract

Long non-coding RNAs (lncRNAs) emerge as critical regulators with various biological functions in living organisms. However, to date, no systematic characterization of lncRNAs has been investigated in the ectoparasitic mite *Varroa destructor*, the most severe biotic threat to honey bees worldwide. Here, we performed an initial genome-wide identification of lncRNAs in *V. destructor* via high-throughput sequencing technology and reported, for the first time, the transcriptomic landscape of lncRNAs in the devastating parasite. By means of a lncRNA identification pipeline, 6,645 novel lncRNA transcripts, encoded by 3,897 gene loci, were identified, including 2,066 sense lncRNAs, 2,772 lincRNAs, and 1,807 lncNATs. Compared with protein-coding mRNAs, *V. destructor* lncRNAs are shorter in terms of full length, as well as of the ORF length, contain less exons, and express at lower level. GO term and KEGG pathway enrichment analyses of the lncRNA target genes demonstrated that these predicted lncRNAs are likely to play key roles in cellular processes, genetic information processing and environmental responses. To our knowledge, this is the first catalog of lncRNA profile in the parasitiformes species, providing a valuable resource for genetic and genomic studies. Understanding the characteristics and features of lncRNAs in *V. destructor* would promote sustainable pest control.

## 1. Introduction

Honey bees, the crucial pollinators all over the world, play a critical role in the balance of the ecosystem, sustainable agriculture and food security [1]. In recent years, however, losses of the western honey bee *Apis mellifera* colonies are a serious issue and there has been a consensus that an emerging ectoparasitic mite, *Varroa destructor*, is the principal threatening factor [2–4]. The ubiquitous parasite originally parasitized in the eastern honey bee *Apis cerana* and shifted host to *A. mellifera* when the latter was introduced into East Asia, the natural distribution range of the former [5,6]. As a result of insufficient coevolution, *V. destructor* presents a tremendous threat to *A. mellifera* apiculture worldwide. In the wake of the occurrence of *V. destructor* in New Zealand in 2000 [7,8] and in Hawaii in 2007 [9], this mite has been distributed globally except for Australia [10].

*V. destructor* feasts on the hemolymph (or fat body) [11,12] of honey bees, transmits viruses and affects host immunity [13–15], severely interrupting the social organization and demographic continuity in *A. mellifera* colonies. Without treatment against this mite, infested *A. mellifera* colonies usually die within six months to two years [16,17]. The parasite lives entirely on its host and cannot survive independently [18] with two life cycle: the phoretic (non-reproductive) phase on the body surface of adult bees and the reproductive phase in the sealed brood cells with immature bees [19]. *V. destructor* reproduction starts from the oogenesis process, which occurs since approximately six hours later after the invaded cell was capped and is crucial for understanding the reproductive biology of the parasite [20,21]. A great number of studies have been performed on this obligate bee parasite, however, molecular studies are very limited as a result of the lack of genomic information. Cornman et al. analyzed *V. destructor* genome sequence, which largely facilitated molecular studies on this mite [22]. Mondet et al. investigated a full life cycle transcriptomic profiling in adult *V. destructor*, capturing the dynamic changes in the mite gene expression [23].

Numerous genome-wide transcriptome has observed that the majority of transcripts do not code for proteins, and these transcripts are referred to as noncoding RNAs (ncRNA) [24]. NcRNA is a generic term for all functional RNAs that are transcribed from DNA but not translated into proteins, including microRNAs (miRNAs), circular RNAs (circRNAs), long non-coding RNAs (lncRNAs), Piwi-interacting RNAs (piRNAs), small interfering RNA (siRNAs), small nucleolar RNAs (snoRNAs), etc. The widespread expression of ncRNAs in eukaryotic cells not only regulates the developmental and physiological functions, but also plays an important role in the process of disease resistance [25]. LncRNAs, most of which are located in the nucleus of eukaryotes, are a cluster of ncRNAs with a length of more than 200 nt, with cap-structure and ploy (A)-tail but usually without a long reading frame [26]. LncRNAs can be classified into four groups based on their positional information on genomes, i.e. sense lncRNAs, intergenic lncRNAs (lincRNAs), intronic lncRNAs (ilncRNAs) and antisense lncRNAs (lncNATs) [27,28]. As functional elements, lncRNAs have been proved to exert their bioactivities by regulating gene expression at the level of epigenetic inheritance, transcription and post transcription, as well as by affecting protein localization and telomere replication [29,30]. Currently, studies of lncRNAs in the field of honey bee science, are however still in its infancy.

The development of the genome-wide transcriptome sequencing technology has enabled novel lncRNA detection. The present a few lncRNA studies in honey bees have demonstrated that these functional elements participate in the regulation of the physiological processes, such as labor division, ovary development, neural networks and pathogen resistance [31–36]. Guo et al. screened the lncRNAs in two honey bee fungi, *Ascospheara apis* and *Nosema ceranae*, unveiling lncRNA studies on honey bee pathogenic agents [37,38]. In the present study, we deeply sequenced the ubiquitous ectoparasite *V. destructor* on the Illumina platform during the oogenesis stage. We identified 6,645 novel lncRNA transcripts corresponding to 3,897 lncRNA genes in the detrimental mite. lncRNAs execute their specified functions with distinct subcellular localization patterns, and most of the lncRNAs were predicted to be located in nucleus and cytoplasm. The structural features and the subcellular localization of the lncRNAs identified in this study showed consistent with their counterparts in the mammals. In order to comprehensively investigate the main biological properties and functions of the target genes of the putative novel lncRNAs, we performed Gene Ontology (GO) and pathway enrichment analyses. Our data offer novel insight into understanding the basic molecular biology of this ubiquitous ectoparasitic mite of honey bees.

## 2. Materials and Methods

### 2.1. Gravid adult female *Varroa destructor* mite collection

*A. mellifera* colonies, located in the apiary of Yangzhou University, Southeast China, kept untreated for half a year were served as *V. destructor* donors. Gravid adult female mites were collected from sealed worker brood cells with the help of tweezers and paint brushes (Figure 1). To obtain the mites with similar physiological status, we experimentally infested the freshly capped worker cells after two-day phoretic phase mimic (see details in [39]). Fifteen adult mites were gathered, two days later after infestation, from the cells as a sample and three samples were prepared. All the mites were frozen in liquid nitrogen after bathing in the phosphate buffered saline, pH 7.4 (Sigma, MO, USA) twice. The samples were stored in −80°C until RNA extraction.

**Figure 1.**
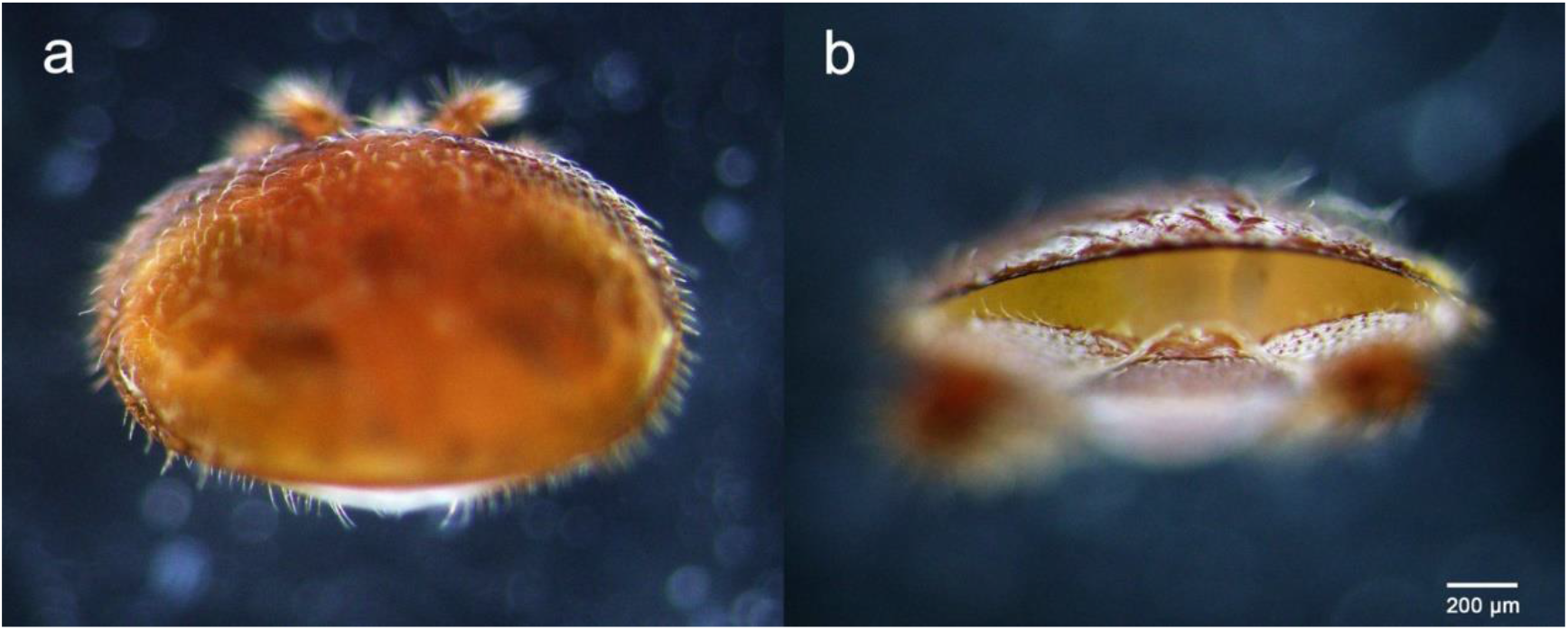
The dorsal (**a**) and post-abdomen (**b**) appearances of the gravid adult female *Varroa destructor* we collected.

### 2.2. Library preparation for lncRNA sequencing

Total RNA was extracted using TRIzol reagent kit (Invitrogen, CA, USA) according to the manufacturer’s protocol. Purity and quantity of total RNA were measured using NanoDrop™ 2000 (Thermo Fisher Scientific, DE, USA). RNA quality was assessed using the RNA Nano 6000 Assay Kit of the Bioanalyzer 2100 system (Agilent Technologies, CA, USA) and was checked on RNase free agarose gel electrophoresis. For the sample preparation, 3 μg RNA was used for each *V. destructor* sample, and NEBNext® UltraTM RNA Library Prep Kit for Illumina® (NEB, MA, USA) was used to generate sequencing libraries. After removing rRNAs, the enriched mRNAs and ncRNAs were fragmented into short fragments by using fragmentation buffer and reverse transcribed into cDNA with random primers. Second-strand cDNA were subsequently synthesized by using DNA polymerase I, RNase H, dNTP and buffer. The cDNA fragments were then purified with QiaQuick PCR extraction kit (Qiagen, Venlo, The Netherlands), end repaired, poly(A) added, and ligated to Illumina sequencing adapters. Uracil-N-Glycosylase was used to digest the second-strand cDNA and the digested products were size selected by agarose gel electrophoresis, PCR amplified, and sequenced on an Illumina HiSeq™ 2500 platform at the Novogene Bioinformatics Institute (Beijing, China). The raw sequencing data were uploaded to the National Centre for Biotechnology Information (SRA accession: PRJNA628859).

### 2.3. LncRNA identification and analyses

Clean reads were obtained by removing reads containing adapter, or ploy-N and low-quality reads from raw data. Meanwhile, we calculated the Q20, Q30 and GC content. The paired-end clean reads were mapped to the reference genome of *V. destructor* (https://www.ncbi.nlm.nih.gov/assembly/GCF_002443255.1) using HISAT2 [40]. We then reconstruct the transcripts with StringTie and HISAT2. To identify new transcripts, all the reconstructed transcripts were aligned to the reference genome and were divided into twelve categories by employing Cuffcompare, and transcripts with one of the five class codes “i”, “j”, “o”, “u” and “x” were potentially recognized as novel ones. The putative novel transcripts were further eliminated by removing the ones with length ≤ 200 nt or with exon number < 2. Coding Potential Calculator (CPC), Coding-Non-Coding Index (CNCI) and Pfam-scan (PFAM) were jointly used to assess the protein-coding potential of the selected novel transcript candidates and the intersection of the results of each software was considered as the candidate set of lncRNAs. We used StringTie again to quantify transcripts abundances by calculating the FPKM (expected fragments per kilobase of transcript per million fragments mapped) values. All the putative novel lncRNAs were computed their subcellular localization by means of lncLocator, an online software for lncRNA location prediction based on a stacked ensemble classifier [41]. Then, we searched coding genes 100 kb upstream and downstream of the predicted novel lncRNAs as *cis* target genes, which were subjected to enrichment analysis of GO functions and KEGG pathways.

The statistical analyses (Student’s *t*-test) of the characteristic differences between lncRNAs and mRNAs were performed with SPSS Statistics 25.

### 2.4. RT-PCR validation

Synthesis of cDNA was performed with RNA products following the manufacturer’s instructions of ReverTra Ace qPCR RT Master Mix (Tiangen, Beijing, China). To validate the putative lncRNAs in *V. destructor* mites, 16 lncRNAs were randomly selected to determine with PCR amplification, which was carried out with the obtained cDNA in a 20 μL reaction volume mixture (2×Taq PCR StarMix; GenStar, Beijing, China) on an Eppendorf cycler. PCR profile consists of a pre-denaturation at 94°C for 5 min; followed by 30 cycles including 94°C for 50 s, 55°C for 30 s and 72°C for 1 min; and a final elongation step at 72°C for 10 min [37]. The tested lncRNAs with their forward and reverse primers were presented in Table S1. PCR products were electrophoresed in 2.5% Tris acetate-EDTA-agarose gel containing 0.01% Gelview (BioTeke, Beijing, China) and visualized under ultraviolet light (Peiqing, Shanghai, China).

## 3. Results

Total RNA of three *V. destructor* pooled samples (Vd-1, Vd-2 and Vd-3) were isolated and sequenced. Overall, 42.8 G sequencing data were generated, corresponding to 285.1 million raw reads and 280.5 quality filtered (clean) reads were generated from the three cDNA libraries (Table 1). G20, G30 and GC content were also shown in Table 1 with the mean values of 97.4%, 92.7% and 43.5%. For the three *V. destructor* samples, 93.2%, 93.6% and 90.0% obtained reads were mapped to the reference genome sequence, and the mapped regions of each sample on the genome were shown in Figure S1. Then 6,645 putative novel non-coding transcripts were predicted using CPC, CNCI and PFAM (Figure 2a), of which 2,066 (31.1%) were sense lncRNAs, 2,772 (41.7%) were lincRNAs, and 1,807 (27.2%) were lncNATs (Figure 2b). IlncRNAs were not observed in this mite. In addition, 32,415 protein coding transcripts were obtained, with 32,331 mapped to the reference genome, and the remaining 84 ones were not annotated.

**Table 1.**
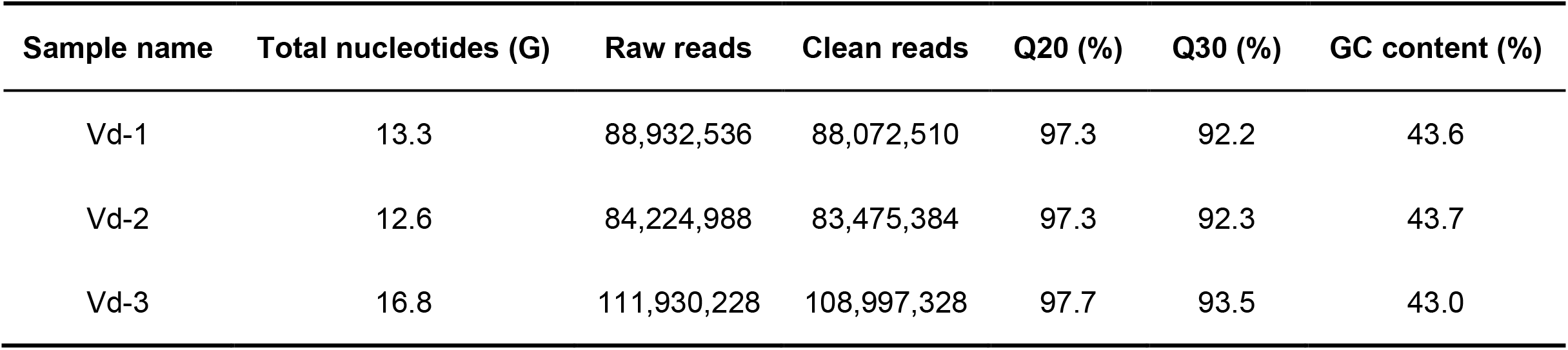
Throughput and quality of RNA-seq of the three libraries.

**Figure 2.**
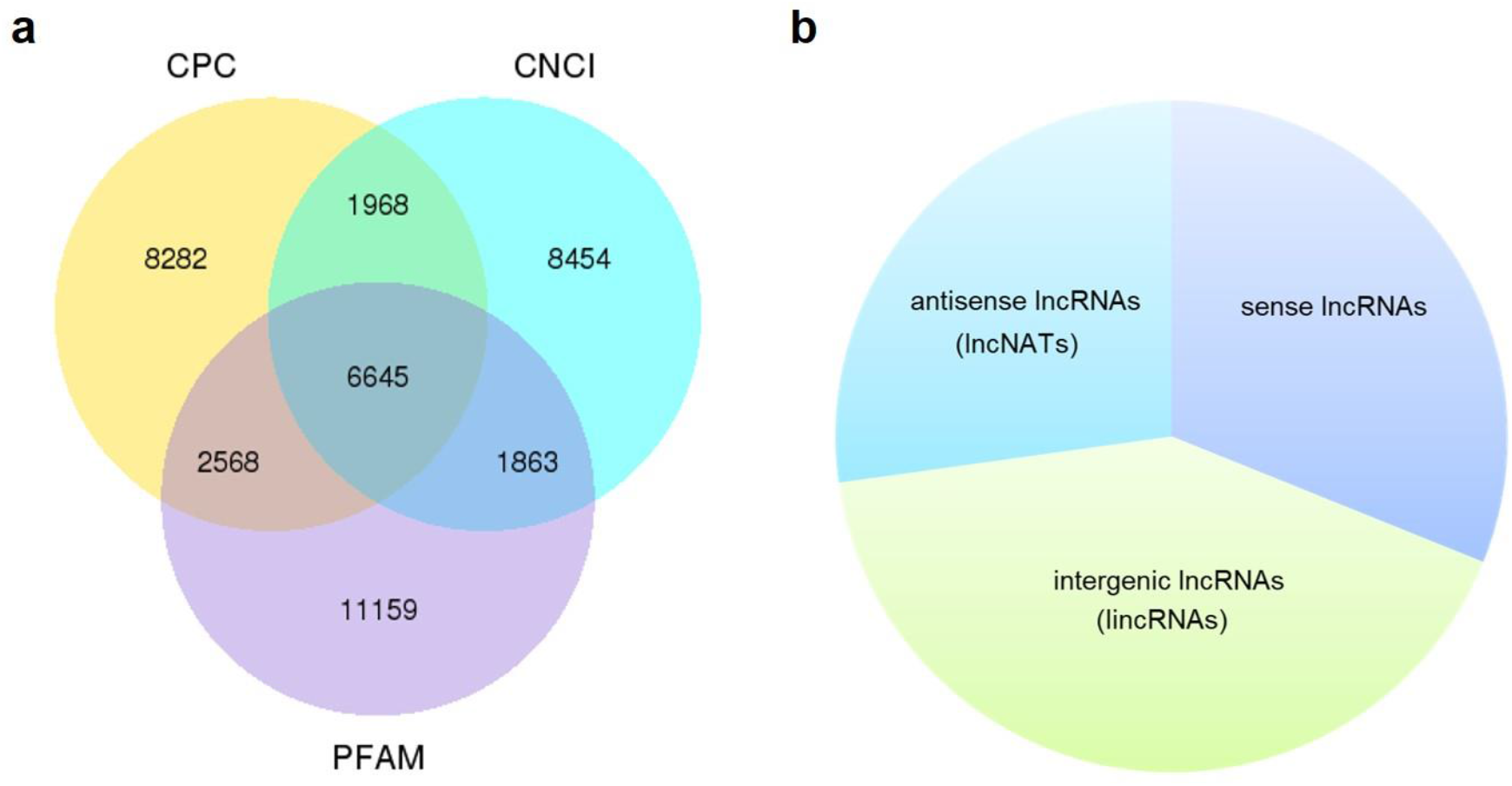
Prediction of novel lncRNA of *V. destructor*. (**a**) Venn analysis of the putative novel lncRNAs by using CPC, CNCI and PFAM. (**b**) The source and distribution percentage of the novel lncRNAs based on the intersection of Venn diagram.

Most lncRNAs contained two exons (55.8%), followed by three (23.8%), four (9.6%), five (4.2%), six (2.2%) and seven (1.1%) (Figure 3a). The ratio of lncRNAs is less than one when the number of exons is greater than seven, and they contained at most 27 exons (Figure 3a). This was significantly different from the coding transcripts (*p* < 0.001), which peaked at six exons (9.5%) and were up to 68 exons. Meanwhile, the ratio of mRNAs with three to eight exons were respectively more than seven (Figure 3a). Most of both lncRNAs (55.4%) and mRNAs (65.5%) ranged from 1,000 bp ~ 5,000 bp in length. But then, 33.2% lncRNAs were less than 1,000 bp and 24.7% mRNAs were between 5,001 bp and 10,000 bp. For the long sequence (> 10,000), 2.1% lncRNAs and 4.7% mRNAs were occupied. As a result, lncRNAs averaged 2,435 bp in length, which was significantly shorter than protein-coding genes (4,187 bp; *p* < 0.001; Figure 3b). Regarding the length of open reading frames (ORFs) in lncRNAs and mRNAs, we got a similar trend with above. Most of the ORFs of both lncRNAs (58.5%) and mRNAs (51.4%) were in the middle range 100 bp ~ 500 bp in length, followed by 39.9% lncRNAs ≤ 100 bp and 31.9% mRNAs ranging from 501 bp to 1,000 bp. Consistently, 0.2% lncRNAs and 16.3% mRNAs were respectively greater than 1,000 bp, and the mean length of ORFs in lncRNAs was significantly shorter than that of mRNAs (132 bp vs 649 bp; *p* < 0.001; Figure 3c). Further, the expression level of lncRNAs showed significantly lower compared to mRNAs (Figure 3d). Additionally, 16 lncRNAs were randomly chosen to be validated with RT-PCR with 15 (93.8%) successful amplification (Figure S2).

**Figure 3.**
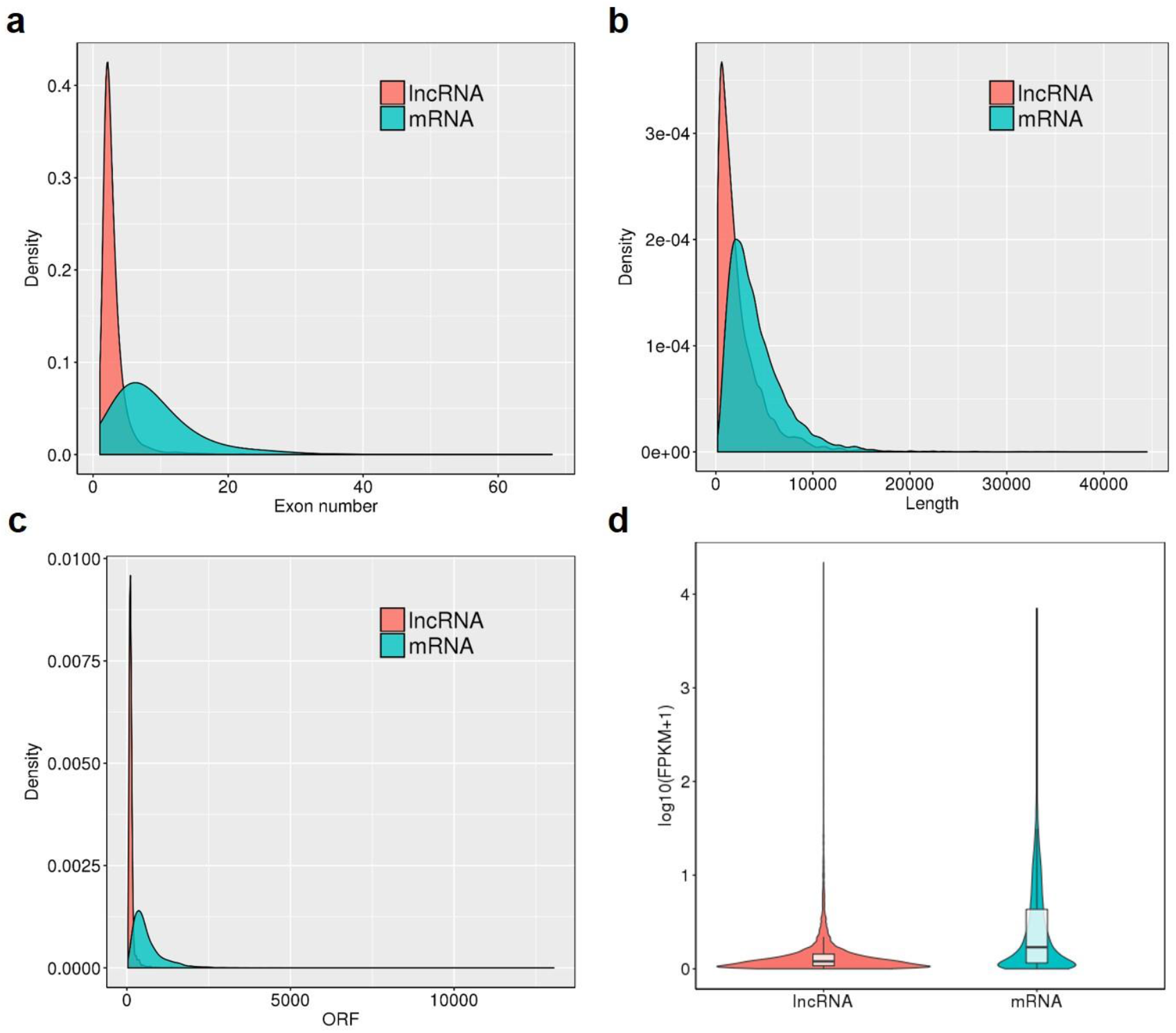
Genomic features of the predicted lncRNAs. Exon number distribution (**a**), length distribution (**b**), ORF length distribution (**c**) and expression level indicated by log10 (FPKM +1) (**d**) of 32,415 coding transcripts (mRNAs) and 6,645 novel lncRNAs were plotted.

As shown in Figure 4, most of the novel lncRNAs were predicted to be localized in the nucleus, followed by in cytoplasm, irrespective of their different sources. In total, only 31, 5 and 5 lncRNAs were predicted in exosome, cytosol and ribosome, respectively. We then obtained 9,500 target genes of novel lncRNAs in *cis* regulation. GO term analysis indicated the target genes were in the ontology class of molecular function and biological process, mainly including protein binding, enzyme activities and metabolic and cellular processes (Figure 5a). The KEGG pathway enrichment indicated that the target genes mostly participated in 104 pathways, which is divided into five classes, i.e. organismal systems, cellular processes, metabolism, environmental information processing and genetic information processing (Figure 5b).

**Figure 4.**
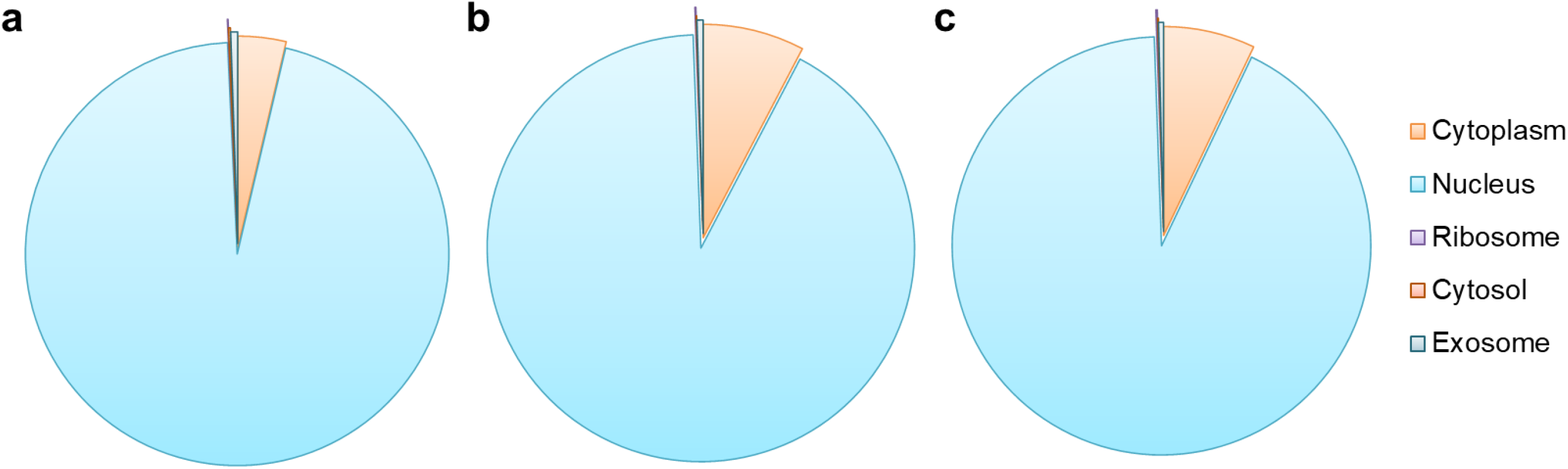
Subcellular localization prediction of the sense lncRNAs (**a**), the intergenic lncRNAs (lincRNAs, **b**) and the antisense lncRNAs (lncNATs, **c**). An online prediction program, lncLocator, which can predict five subcellular localizations of lncRNAs, was used for this analysis.

**Figure 5.**
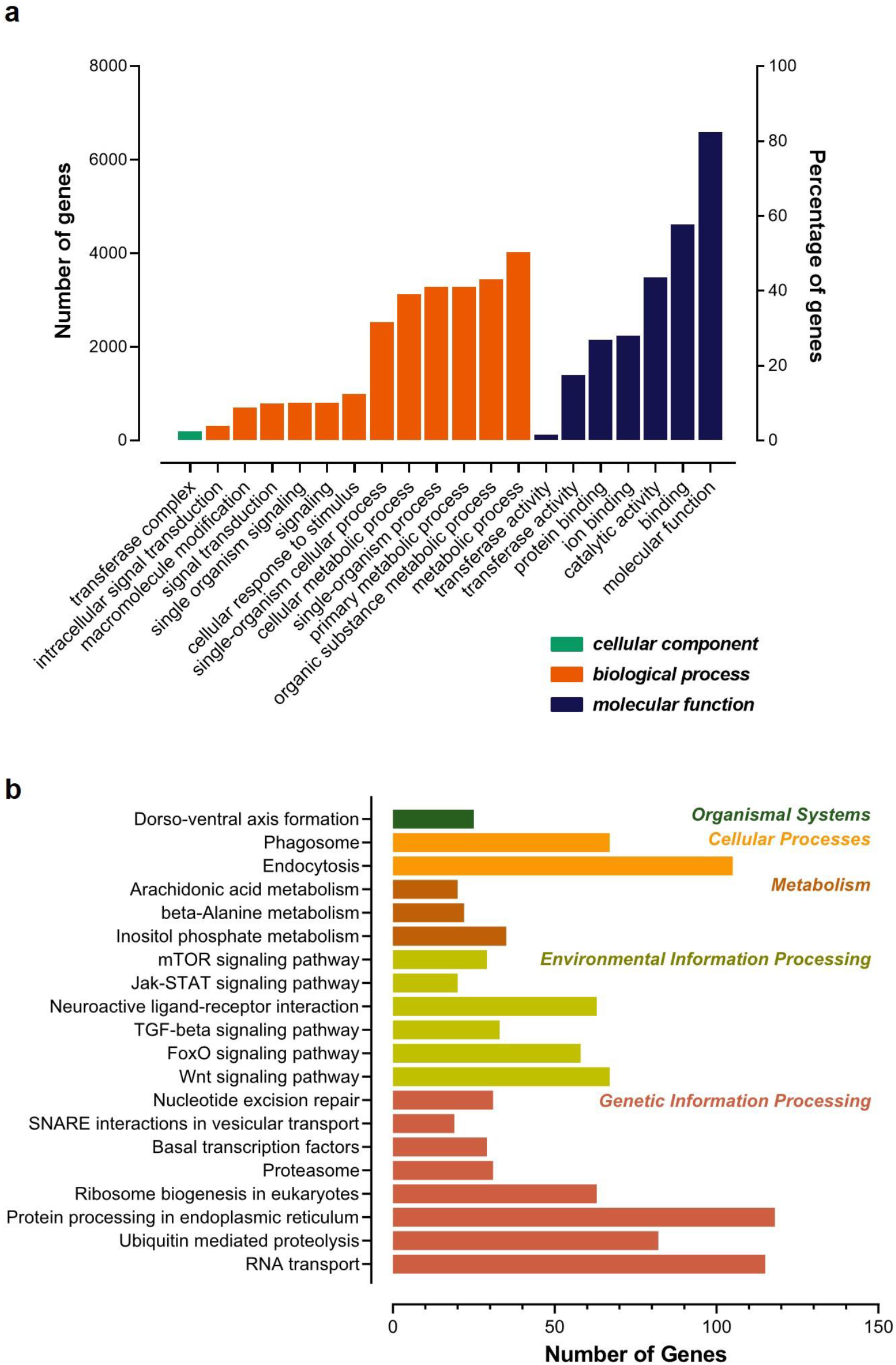
GO categorization (**a**) and KEGG pathway (**b**) analyses for the target genes of the predicted lncRNAs. Top 20 enriched terms were respectively shown.

## 4. Discussion

LncRNAs have emerged as critical participators in a variety of cellular activities, ranging from simple housekeeping to complex regulatory functions. However, till now, the studies of lncRNAs are mainly conducted in the field of humans, mammal and crops. In contrast, the research on invertebrates is still at the early stage. Here, we identified the sequences and expression features of the lncRNAs via high-throughput sequencing technology, for the first time, in a devastating ectoparasite, *V. destructor*, of the chief pollinator, honey bee. We obtained 6,645 putative novel lncRNAs from 3,897 gene loci in the *V. destructor*, including 2,066 sense lncRNAs, 2,772 lincRNAs, and 1,807 lncNATs. The characteristics of the lncRNAs and the comparison with mRNAs were also reported. To verify the reliability of the RNA-seq results and the predicted lncRNAs, 16 non-coding transcripts were randomly selected to do RT-PCR validation, and 93.8% lncRNAs amplified signal bands denoting a reliable output of lncRNAs identified in this study. To the best of our knowledge, lncRNA studies have not performed in the parasitiformes species before.

*V. destructor* reproduction is limited to a short window when the immature honey bee host is concealed in wax cells. Mated female mites target to meal on the larva host five hours after the invaded cell was capped [42,43] and initiates oogenesis about another hour later [21]. The mite then lays the first male egg approximately 70 hours after the cell capping [19,42]. Considering these timing, we collected the mites 48 hours later after the mite was introduced into the freshly capped cells. The artificial infestation has been established to be a suitable method for *V. destructor* research [20,39,44], and mites parasitized in the capped brood cells are less variable in physiology and fitness than on the adult bee bodies [45]. The mites collected were further confirmed gravid with dilated post-abdomen, from which we can even see the eggs inside under microscope (Fig 1b).

Just as lncRNA screening in other species, lincRNAs, the most extensively studied category of lncRNAs, usually account for the largest proportion [37,46–48], although some studies show otherwise [38]. We did not detect any ilncRNAs, which were regarded as the lowest conservative class of lncRNAs [49], in *V. destructor*. Intriguingly, this is also the case of *N. ceranae*, another pathogenic agent of honey bees [38]. Most of the *V. destructor* lncRNAs contained two exons with the average of 3.0 exons, significantly less than mRNAs, which was also in line with the related studies of lncRNAs on other invertebrate counterparts, even mammalian and plant species [36–38,47,48,50,51]. Besides, the shorter length of the putative lncRNAs and the ORFs compared to mRNA also shared similar features with other well-studied species [36–38,47–51]. Although lncRNAs always performed lower expression level than the protein-coding genes [this study, 31,36–38], their role in the functional activities of organisms has been widely proved to be of significance [29–31,36,47,51].

LncRNAs can function in *cis* to regulate the transcriptional expression of neighboring genes on the same allele [52]. The upstream lncRNAs with intersection of promoter or other *cis* elements may regulate gene expression at the level of transcription or post-transcription, and lncRNAs in the downstream or 3’UTR region may have other regulatory functions. LncRNAs in less than 100 kb up/down stream of a gene may serve as *cis* regulatory factors. The *cis* target genes were engaged in various functions associated with diverse molecular functions and biological processes (Figure 5a). The top three enriched pathways were protein processing in endoplasmic reticulum, RNA transport and endocytosis, and each of them were enriched by more than 100 genes (Figure 5b). KEGG analysis suggests that most of the target genes of the lncRNAs are engaged in the translation, folding, sorting and degradation, transport and catabolism. In addition, the target genes are also included in the genetic and environmental information processing, metabolism, development and regeneration, signaling molecules and interaction and so forth. Notably, 25 genes were enriched in a pathway of dorso-ventral axis formation (Figure 5b), suggesting that they may play a crucial role in the development and regeneration during the process of oogenesis of the mite. Therefore, *V. destructor* lncRNAs may be potentially responsible for the regulatory functions of cellular and biological progress.

Similar to proteins, lncRNAs are of importance to be localized in specific cellular compartments, which provides insights for understanding their complex biological functions [53]. We predicted the subcellular localizations of the identified sense lncRNAs, lincRNAs and lncNATs, of which 95.6%, 91.7% and 92.4% were respectively accumulated in nucleus (Figure 4). These lncRNAs have been proposed to play strong roles in nuclear architecture and gene expression regulation. They are associated with chromatin-modifying complexes, directly influence transcription, act as precursors for small RNAs, participate in stem cell pluripotency and differentiation and so forth [54]. Cytoplasmic lncRNAs, the second most popularly located lncRNAs, have been evidenced to impact gene expression in a variety of ways, such as interfering with protein post-translational modifications with a result of aberrant signal transduction [55], acting as decoys for miRNAs and proteins [56,57] and affecting mRNA translation in the cytoplasm [58]. Hence, the lncRNAs in distinct subcellular compartments are of great interest to decipher their diverse functional significance.

## 5. Conclusions

*V. destructor* presenting tremendous threat to apiculture worldwide, the in-depth molecular studies on the parasite will facilitate to control this unpopular pest. We reported the lncRNA profile of *V. destructor* by genome-wide RNA-seq in this study, and the genomic and structural features of the lncRNAs showed consistent with their counterparts in other species. Evidence is becoming increasingly clear that the function of lncRNAs is associated with their unique subcellular localization, and most of the lncRNAs detected in *V. destructor* were accumulated in nucleus and cytoplasm. The target genes of lncRNAs were inferred to participate in diverse regulatory functions via *cis* regulation by GO term and KEGG pathway enrichment analyses. Our data provide genetic resources for exploration of the functional roles of lncRNAs involved in the ectoparasite *V. destructor*. Further studies would be of interest and value to characterize the expression profile of lncRNAs in the different life stages of the ubiquitous mite.

## Author Contributions

Conceptualization, T.J., G.C. and Z.L.; methodology, T.J., X.S. and Z.L.; validation, Y.L., X.C. and C.H.; formal analysis, Z.L., Y.L., C.H. and X.S.; investigation, Y.L., X.C., H.C. and H.X.; resources, W.W. and Y.K.; data curation, X.X.; writing—original draft preparation, Z.L.; writing—review and editing, X.S., W.W., Y.K. and T.J.; supervision, T.J. and G.C.; project administration, T.J. and G.C.; funding acquisition, T.J., G.C. and Z.L.

## Supplementary Materials

**Table S1.** Primer sets used for RT-PCR validation of the identified lncRNAs.

**Figure S1.** Distribution percentage of the clean reads from Vd-1(**a**), Vd-2(**b**) and Vd-3(**c**) mapped to the genome regions of *Varroa destructor*.

**Figure S2.** RT-PCR validation of the randomly selected 16 lncRNAs. DNA marker (M) were used to indicate the product size in the left lane. Lane 1 to lane 16 were 16 lncRNAs orderly listed in the Table S1.

## References

1. Potts, S.G.; Imperatriz-Fonseca, V.; Ngo, H.T.; Aizen, M.A.; Biesmeijer, J.C.; Breeze, T.D.; Dicks, L.V.; Garibaldi, L.A.; Hill, R.; Settele, J., et al. Safeguarding pollinators and their values to human well-being. Nature 2016, 540, 220–229, doi:10.1038/nature20588.

2. Evans, J.D. Selection for barriers between honey bees and a devastating parasite. Mol. Ecol. 2019, 28, 2955–2957, doi:10.1111/mec.15142.

3. Nazzi, F.; Conte, Y.L. Ecology of *Varroa destructor*, the Major Ectoparasite of the Western Honey Bee, *Apis mellifera*. Annu. Rev. Entomol. 2016, 61, 417–432, doi:10.1146/ANNUREV-ENTO-010715-023731.

4. Neumann, P.; Carreck, N.L. Honey bee colony losses. J. Apic. Res. 2010, 49, 1–6, doi:10.3896/IBRA.1.49.1.01.

5. Akratanakul, P.; Burgett, M. *Varroa jacobsoni*: A prospective pest of honeybees in many parts of the world. Bee World 1975, 56, 119–121, doi:10.1080/0005772X.1975.11097554.

6. Oldroyd, B.P. Coevolution while you wait: *Varroa jacobsoni*, a new parasite of Western honeybees. Trends Ecol. Evol. 1999, 14, 312–315, doi:10.1016/S0169-5347(99)01613-4.

7. Mondet, F.; de Miranda, J.R.; Kretzschmar, A.; Le Conte, Y.; Mercer, A.R. On the front line: Quantitative virus dynamics in honeybee (*Apis mellifera L.*) colonies along a new expansion front of the parasite *Varroa destructor*. PLoS Pathog. 2014, 10, e1004323, doi:10.1371/journal.ppat.1004323.

8. Todd, J.H.; Miranda, J.R.D.; Ball, B.V. Incidence and molecular characterization of viruses found in dying New Zealand honey bee (*Apis mellifera*) colonies infested with *Varroa destructor*. Apidologie 2007, 38, 354–367, doi:10.1051/APIDO:2007021.

9. Martin, S.J.; Highfield, A.C.; Brettell, L.; Villalobos, E.M.; Budge, G.E.; Powell, M.; Nikaido, S.; Schroeder, D.C. Global honey bee viral landscape altered by a parasitic mite. Science 2012, 336, 1304–1306, doi:10.1126/science.1220941.

10. Genersch, E. Honey bee pathology: current threats to honey bees and beekeeping. Appl. Microbiol. Biotechnol. 2010, 87, 87–97, doi:10.1007/s00253-010-2573-8.

11. Ramsey, S.; Gulbronson, C.; Mowery, J.; Ochoa, R.; VanEngelsdorp, D.; Bauchan, G. A multi-microscopy approach to discover the feeding site and host tissue consumed by *Varroa destructor* on host honey bees. Microsc. Microanal. 2018, 24, 1258–1259, doi:10.1017/S1431927618006773.

12. Ramsey, S.D.; Ochoa, R.; Bauchan, G.; Gulbronson, C.; Mowery, J.D.; Cohen, A.; Lim, D.; Joklik, J.; Cicero, J.M.; Ellis, J.D., et al. Varroa destructor feeds primarily on honey bee fat body tissue and not hemolymph. Proc. Natl. Acad. Sci. U. S. A. 2019, 116, 1792–1801, doi:10.1073/pnas.1818371116.

13. Di Prisco, G.; Annoscia, D.; Margiotta, M.; Ferrara, R.; Varricchio, P.; Zanni, V.; Caprio, E.; Nazzi, F.; Pennacchio, F. A mutualistic symbiosis between a parasitic mite and a pathogenic virus undermines honey bee immunity and health. Proc. Natl. Acad. Sci. U. S. A. 2016, 113, 3203–3208, doi:10.1073/pnas.1523515113.

14. Yang, X.; Cox-Foster, D.L. Impact of an ectoparasite on the immunity and pathology of an invertebrate: Evidence for host immunosuppression and viral amplification. Proc. Natl. Acad. Sci. U. S. A. 2005, 102, 7470–7475, doi:10.1073/pnas.0501860102.

15. Rosenkranz, P.; Aumeier, P.; Ziegelmann, B. Biology and control of *Varroa destructor*. J. Invertebr. Pathol. 2010, 103, S96–S119.

16. Kraus, B.; Page, R.E. Effect of *Varroa jacobsoni* (Mesostigmata: Varroidae) on feral *Apis mellifera* (Hymenoptera: Apidae) in California. Environ. Entomol. 1995, 24, 1473–1480, doi:10.1093/EE/24.6.1473.

17. Le Conte, Y.; Ellis, M.; Ritter, W. *Varroa mites* and honey bee health: can *Varroa* explain part of the colony losses? Apidologie 2010, 41, 353–363, doi:10.1051/apido/2010017.

18. Traynor, K.; Mondet, F.; Miranda, J.d.; Techer, M.; Kowallik, V.; Oddie, M.; Chantawannakul, P.; McAfee, A. *Varroa destructor*: A complex parasite, crippling honeybees worldwide. 2020; 10.20944/PREPRINTS202002.0374.V1.

19. Martin, S.J. Ontogenesis of the mite *Varroa jacobsoni* Oud. in worker brood of the honeybee *Apis mellifera L*. under natural conditions. Exp. Appl. Acarol. 1994, 19, 199–210, doi:10.1007/BF00130823.

20. Häußermann, C.K.; Giacobino, A.; Munz, R.; Ziegelmann, B.; Palacio, M.A.; Rosenkranz, P. Reproductive parameters of female *Varroa destructor* and the impact of mating in worker brood of *Apis mellifera*. Apidologie 2019, doi:10.1007/S13592-019-00713-9.

21. Garrido, C.; Rosenkranz, P.; Stürmer, M.; Rübsam, R.; Büning, J. Toluidine blue staining as a rapid measure for initiation of oocyte growth and fertility in *Varroa jacobsoni* Oud. Apidologie 2000, 31, 559–566, doi:10.1051/APIDO:2000146.

22. Cornman, S.R.; Schatz, M.C.; Johnston, S.J.; Chen, Y.p.; Pettis, J.; Hunt, G.; Bourgeois, L.; Elsik, C.; Anderson, D.; Grozinger, C.M., et al. Genomic survey of the ectoparasitic mite *Varroa destructor*, a major pest of the honey bee *Apis mellifera*. BMC Genomics 2010, 11, 602–602, doi:10.1186/1471-2164-11-602.

23. Mondet, F.; Rau, A.; Klopp, C.; Rohmer, M.; Severac, D.; Le Conte, Y.; Alaux, C. Transcriptome profiling of the honeybee parasite *Varroa destructor* provides new biological insights into the mite adult life cycle. BMC Genomics 2018, 19, 328, doi:10.1186/s12864-018-4668-z.

24. Lee, T.-L.; Xiao, A.; Rennert, O.M. Identification of novel long noncoding RNA transcripts in male germ cells. In Germline Development: Methods and Protocols, Chan, W.-Y., Blomberg, L.A., Eds. Springer New York: New York, NY, 2012; 10.1007/978-1-61779-436-0_9pp. 105–114.

25. Esteller, M. Non-coding RNAs in human disease. Nat. Rev. Genet. 2011, 12, 861–874, doi:10.1038/nrg3074.

26. Guttman, M.; Amit, I.; Garber, M.; French, C.; Lin, M.F.; Feldser, D.; Huarte, M.; Zuk, O.; Carey, B.W.; Cassady, J.P., et al. Chromatin signature reveals over a thousand highly conserved large non-coding RNAs in mammals. Nature 2009, 458, 223–227, doi:10.1038/nature07672.

27. Harrow, J.; Frankish, A.; Gonzalez, J.M.; Tapanari, E.; Diekhans, M.; Kokocinski, F.; Aken, B.L.; Barrell, D.; Zadissa, A.; Searle, S., et al. GENCODE: The reference human genome annotation for The ENCODE Project. Genome Res. 2012, 22, 1760–1774, doi:10.1101/GR.135350.111.

28. Ma, L.; Bajic, V.B.; Zhang, Z. On the classification of long non-coding RNAs. RNA Biol. 2013, 10, 924–933, doi:10.4161/RNA.24604.

29. Furuno, M.; Pang, K.C.; Ninomiya, N.; Fukuda, S.; Frith, M.C.; Bult, C.; Kai, C.; Kawai, J.; Carninci, P.; Hayashizaki, Y., et al. Clusters of internally primed transcripts reveal novel long noncoding RNAs. PLoS Genet. 2006, 2, 537–553, doi:10.1371/JOURNAL.PGEN.0020037.

30. Mercer, T.R.; Dinger, M.E.; Mattick, J.S. Long non-coding RNAs: insights into functions. Nat. Rev. Genet. 2009, 10, 155–159, doi:10.1038/NRG2521.

31. Chen, D.; Chen, H.; Du, Y.; Zhou, D.; Geng, S.; Wang, H.; Wan, J.; Xiong, C.; Zheng, Y.; Guo, R. Genome-wide identification of long non-coding RNAs and their regulatory networks lnvolved in *Apis mellifera ligustica* response to *Nosema ceranae* infection. Insects 2019, 10, doi:10.3390/INSECTS10080245.

32. Chen, X.; Ma, C.; Chen, C.; Lu, Q.; Shi, W.; Liu, Z.; Wang, H.; Guo, H. Integration of lncRNA–miRNA–mRNA reveals novel insights into oviposition regulation in honey bees. PeerJ 2017, 5, e3881, doi:10.7717/peerj.3881.

33. Humann, F.C.; Tiberio, G.J.; Hartfelder, K. Sequence and expression characteristics of long noncoding RNAs in honey bee caste development – potential novel regulators for transgressive ovary size. PLoS One 2013, 8, e78915, doi:10.1371/journal.pone.0078915.

34. Jayakodi, M.; Jung, J.W.; Park, D.; Ahn, Y.-J.; Lee, S.-C.; Shin, S.-Y.; Shin, C.; Yang, T.-J.; Kwon, H.W. Genome-wide characterization of long intergenic non-coding RNAs (lincRNAs) provides new insight into viral diseases in honey bees *Apis cerana* and *Apis mellifera*. BMC Genomics 2015, 16, 680, doi:10.1186/s12864-015-1868-7.

35. Kiya, T.; Kunieda, T.; Kubo, T. Inducible- and constitutive-type transcript variants of kakusei, a novel non-coding immediate early gene, in the honeybee brain. Insect Mol. Biol. 2008, 17, 531–536, doi:10.1111/j.1365-2583.2008.00821.x.

36. Liu, F.; Shi, T.; Qi, L.; Su, X.; Wang, D.; Dong, J.; Huang, Z.Y. lncRNA profile of *Apis mellifera* and its possible role in behavioural transition from nurses to foragers. BMC Genomics 2019, 20, 393, doi:10.1186/s12864-019-5664-7.

37. Guo, R.; Chen, D.; Xiong, C.; Hou, C.; Zheng, Y.; Fu, Z.; Diao, Q.; Zhang, L.; Wang, H.; Hou, Z., et al. Identification of long non-coding RNAs in the chalkbrood disease pathogen *Ascospheara apis*. J. Invertebr. Pathol. 2018, 156, 1–5, doi:10.1016/J.JIP.2018.06.001.

38. Guo, R.; Chen, D.; Xiong, C.; Hou, C.; Zheng, Y.; Fu, Z.; Liang, Q.; Diao, Q.; Zhang, L.; Wang, H., et al. First identification of long non-coding RNAs in fungal parasite *Nosema ceranae*. Apidologie 2018, 49, 660–670, doi:10.1007/S13592-018-0593-Z.

39. Lin, Z.; Qin, Y.; Page, P.; Wang, S.; Li, L.; Wen, Z.; Hu, F.; Neumann, P.; Zheng, H.; Dietemann, V. Reproduction of parasitic mites *Varroa* destructor in original and new honeybee hosts. Ecol. Evol. 2018, 8, 2135–2145, doi:10.1002/ece3.3802.

40. Kim, D.; Langmead, B.; Salzberg, S.L. HISAT: A fast spliced aligner with low memory requirements. Nat. Methods 2015, 12, 357–360, doi:10.1038/NMETH.3317.

41. Cao, Z.; Pan, X.; Yang, Y.; Huang, Y.; Shen, H.-B. The lncLocator: a subcellular localization predictor for long non-coding RNAs based on a stacked ensemble classifier. Bioinformatics2018, 34, 2185–2194, doi:10.1093/bioinformatics/bty085.

42. Donzé, G.; Guerin, P.M. Behavioral attributes and parental care of *Varroa* mites parasitizing honeybee brood. Behav. Ecol. Sociobiol. 1994, 34, 305–319, doi:10.1007/BF00197001.

43. Ifantidis, M.D. Some aspects of the process of *Varroa jacobsoni* mite entrance into honey bee (*Apis mellifera*) brood cells. Apidologie 1988, 19, 387–396, doi:10.1051/APIDO:19880406.

44. Dietemann, V.; Nazzi, F.; Martin, S.J.; Anderson, D.L.; Locke, B.; Delaplane, K.S.; Wauquiez, Q.; Tannahill, C.; Frey, E.; Ziegelmann, B., et al. Standard methods for varroa research. J. Apic. Res. 2013, 52, 1–54, doi:10.3896/IBRA.1.52.1.09.

45. Milani, N. The resistance of *Varroa jacobsoni* Oud to pyrethroids: a laboratory assay. Apidologie 1995, 26, 415–429, doi:10.1051/APIDO:19950507.

46. Sun, J.; Lin, Y.; Wu, J. Long non-coding RNA expression profiling of mouse testis during postnatal development. PLoS One 2013, 8, e75750, doi:10.1371/journal.pone.0075750.

47. Wu, Y.; Cheng, T.; Liu, C.; Liu, D.; Zhang, Q.; Long, R.; Zhao, P.; Xia, Q. Systematic Identification and characterization of long non-coding RNAs in the Silkworm, Bombyx mori. PLoS One 2016, 11, e0147147, doi:10.1371/journal.pone.0147147.

48. Zhan, S.; Dong, Y.; Zhao, W.; Guo, J.; Zhong, T.; Wang, L.; Li, L.; Zhang, H. Genome-wide identification and characterization of long non-coding RNAs in developmental skeletal muscle of fetal goat. BMC Genomics 2016, 17, 666, doi:10.1186/s12864-016-3009-3.

49. Wang, Y.; Xue, S.; Liu, X.; Liu, H.; Hu, T.; Qiu, X.; Zhang, J.; Lei, M. Analyses of long non-coding RNA and mRNA profiling using RNA sequencing during the pre-implantation phases in pig endometrium. Sci. Rep. 2016, 6, 20238, doi:10.1038/srep20238.

50. Trapnell, C.; Williams, B.A.; Pertea, G.; Mortazavi, A.; Kwan, G.; van Baren, M.J.; Salzberg, S.L.; Wold, B.J.; Pachter, L. Transcript assembly and quantification by RNA-Seq reveals unannotated transcripts and isoform switching during cell differentiation. Nat. Biotechnol. 2010, 28, 511–515, doi:10.1038/nbt.1621.

51. Zhang, Y.-C.; Liao, J.-Y.; Li, Z.-Y.; Yu, Y.; Zhang, J.-P.; Li, Q.-F.; Qu, L.-H.; Shu, W.-S.; Chen, Y.-Q. Genome-wide screening and functional analysis identify a large number of long noncoding RNAs involved in the sexual reproduction of rice. Genome Biol. 2014, 15, 512, doi:10.1186/s13059-014-0512-1.

52. Robinson, E.K.; Covarrubias, S.; Carpenter, S. The how and why of lncRNA function: An innate immune perspective. Biochim. Biophys. Acta, Gene Regul. Mech. 2020, 1863, 194419, doi:10.1016/j.bbagrm.2019.194419.

53. Chen, L.-L. Linking long noncoding RNA localization and function. Trends Biochem. Sci. 2016, 41, 761–772, doi:10.1016/j.tibs.2016.07.003.

54. Chen, L.-L.; Carmichael, G.G. Decoding the function of nuclear long non-coding RNAs. Curr. Opin. Cell Biol. 2010, 22, 357–364, doi:10.1016/j.ceb.2010.03.003.

55. Lin, A.; Li, C.; Xing, Z.; Hu, Q.; Liang, K.; Han, L.; Wang, C.; Hawke, D.H.; Wang, S.; Zhang, Y., et al. The LINK-A lncRNA activates normoxic HIF1α signalling in triple-negative breast cancer. Nat. Cell Biol. 2016, 18, 213–224, doi:10.1038/ncb3295.

56. Cesana, M.; Cacchiarelli, D.; Legnini, I.; Santini, T.; Sthandier, O.; Chinappi, M.; Tramontano, A.; Bozzoni, I. A long noncoding RNA controls muscle differentiation by functioning as a competing endogenous RNA. Cell 2011, 147, 358–369, doi:10.1016/j.cell.2011.09.028.

57. Lee, S.; Kopp, F.; Chang, T.-C.; Sataluri, A.; Chen, B.; Sivakumar, S.; Yu, H.; Xie, Y.; Mendell, Joshua T. Noncoding RNA NORAD regulates genomic stability by sequestering PUMILIO proteins. Cell 2016, 164, 69–80, doi:10.1016/j.cell.2015.12.017.

58. Gong, C.; Maquat, L.E. lncRNAs transactivate STAU1-mediated mRNA decay by duplexing with 3′ UTRs via Alu elements. Nature 2011, 470, 284–288, doi:10.1038/nature09701.

